# DNA-mediated self-assembly of gold nanoparticles on protein superhelix

**DOI:** 10.1101/449561

**Authors:** Tao Zhang, Ingemar André

## Abstract

Recent advances in protein engineering have enabled methods to control the self-assembly of protein on various length-scales. One attractive application for designed proteins is to direct the spatial arrangement of nanomaterials of interest. Until now, however, a reliable conjugation method is missing to facilitate site-specific positioning. In particular, bare inorganic nanoparticles tend to aggregate in the presence of buffer conditions that are often required for the formation of stable proteins. Here, we demonstrated a DNA mediated conjugation method to link gold nanoparticles with protein structures. To achieve this, we constructed *de novo* designed protein fibers based on previously published uniform alpha-helical units. DNA modification rendered gold nanoparticles with increased stability against ionic solutions and the use of complementary strands hybridization guaranteed the site-specific binding to the protein. The combination of high resolution placement of anchor points in designed protein assemblies with the increased control of covalent attachment through DNA binding can enable investigations of multilevel physical coupling events of nanocomponents on protein templates and expand the application of protein structures to material sciences.

---

Investigations on distance and orientation-dependent interactions among self-assembled nanocomponents can determine manifold physical coupling events on nanoscale or even to reveal new phenomena in low dimensional systems. However, it is far from trivial to observe these interactions due to their small sizes, significant surface effects, and constant thermal fluctuations in solution. One possible solution is to fix the nanocomponents in place *via* bottom-up self-assembly.^1,2^ For example, DNA-based assembly employs the complementarity between DNA base-pair interactions to fold numerous structures with nanometer-scale accuracy in a one-pot reaction.^3–6^ DNA structures carrying docking strands in desired places can be used as templates for site-specific positioning of nanoparticles and the subsequent studies on their physical interactions and collective coupling effects.^7,8^ The last three decades have seen the successful development in DNA nanotechnology. Recent publications^9,10^ provide further discussion of the advancements in DNA-based structuring technology and the use of DNA nanostructures as scaffolds for creating plasmonic architectures.

*De novo* protein design is an attractive alternative to DNA-based assembly for customized construction of nano-objects and can sometimes implement the placement of atoms with close to atomic resolution.^11–14^ Computational protein design is capable of exploring a vast number of sequences and structures *in silico*, far beyond what can be experimentally tested, and access structural states that have not been sampled in evolution.^15–17^ Although *de novo* design is still a challenging prospect, stable protein structures with desired geometries from one dimensional fibres to three dimensional crystals have been successfully designed and verified.^18–21^ Because the position of individual residues is typically well-defined in rationally designed proteins, such proteins are also able to serve as a robust platform for scaffolding the arrangement of guest molecules, with potential applications in sensing, nanophotonics, and as functional entities.

To facilitate the binding of guest components on protein, various of robust chemistry methods^22–26^ are available. However, approaches are still demanded to realize precise assembly at addressable sites, especially when manipulating multiple nanocomponents in parallel. For example, to attach gold nanoparticles on protein structure, pre-synthesized gold nanoparticles can be captured on protein *via* gold-reactive groups such as cysteine,^27,28^ or *via* in situ growth.^29^ In the case of the former, unbound gold nanoparticles tend to form clusters under the buffer conditions necessary to stabilize the protein, while for the latter, care must be taken to ensure the monodispersity for nanoparticles’ shapes and sizes. To address this problem, here we present a method of DNA-mediated gold nanoparticles assembly^30^ and apply the method to label a *de novo* designed protein that forms a helical fibre structure. The protein superhelix was assembled from homogeneous building blocks consisting of six pairs of repeated alpha helix segments from a previously published design.^31^ DNA was selected as a binding partner to link gold nanoparticles and protein structures. Thiol-gold bonding is the standard approach to conjugate thiolated-DNA with gold surfaces and the complementary DNA oligonucleotides linkage to protein structures can be completed *via* amine-to-amine or thiol-to-amine coupling. Thereafter, DNA strands hybridization enables the site-specific assembly of DNA-modified gold nanoparticles on protein templates.

Our protein fibre is based on a repeat protein design composed of helix-loop-helix-loop structural motif introduced by T.J. Brunette *et al*..^31^ Repetition of the helix-loop-helix-loop results in a supramolecular helical shape of the repeat protein. In their constructs, the design consists of four pairs of alpha helix bundles with two inner pairs of repeated helix bundles and two hydrophilic caps at both ends to endow a good solubility of the proteins. We reasoned that a fibrillar structure with helical symmetry could be formed by removing the capping modules. As a basis for this design we selected one of the repeat proteins presented by T.J. Brunette *et al*. because of its favourable twist and radius (Figure 1A, two inner repeated helix bundles highlighted in green, two end caps in cyan, and the hydrophobic interfaces in yellow) and fused three copies of the original unit (Figure 1B) into a monomer containing six pairs of repeated alpha helix bundles (Figure 1C). Linking six pairs together can significantly improve the structure solubility of the construct even without the hydrophilic caps. As the interfaces at the ends of the monomer are no different from the interfaces between each pair of alpha helix, each monomers can be engaged together from plus end (+) to minus end (-) through the interface interactions as designed in the original protein. Due to an offset between the neighbouring alpha-helix pairs in the model protein design, the accumulated long-range assembly therefore generates a helical fibre. Note that plus (+) and minus (-) are labelled to assist the explanation of interfaces engaged self-assembly, but no preferential directions for the fibre growth are expected.

**Figure 1:**
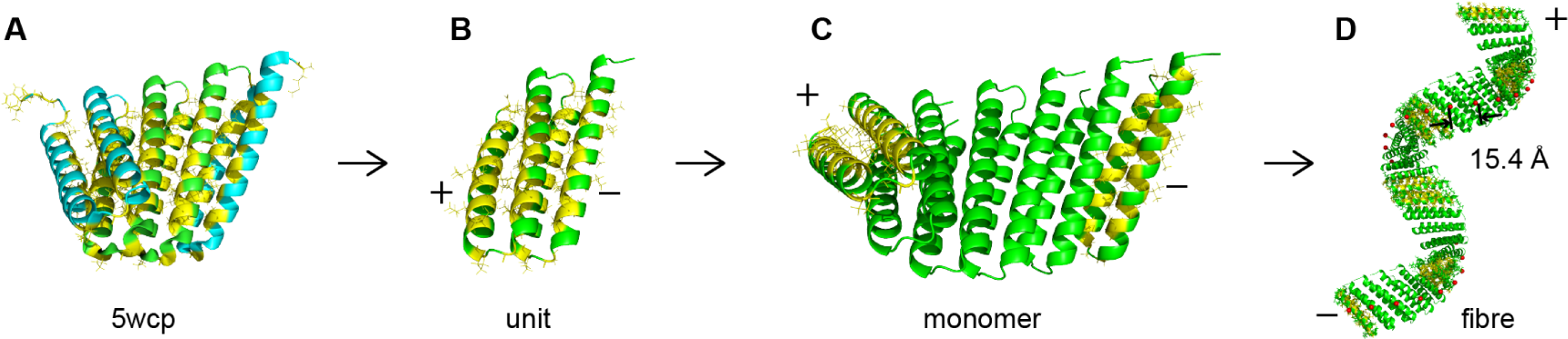
Schematic illustration for generating a protein superhelix. A, the original chiral protein design (PDB 5CWP) is adopted from ref [31]. The two repeated alpha helix pairs are highlighted in green and the two end caps in cyan. Hydrophobic residues at interface regions are marked in yellow. B, the unit is comprised of two repeated alpha helix pairs. The plus and minus labels indicate that the hydrophobic interfaces are active for further assembly, where minus end can be associated with the plus end. Note that there are no preferential directions for the fibre growth. C, the monomer used in the experiments is created by fusing three units (panel B) together. D, the interface engagement between the monomer lead to the assembly of a right handed superhelix. Lysine nitrogen atoms are highlighted with red spheres and there is well defined spacing of 15.4 Å between adjacent lysine groups.

As mentioned above, protein labelling requires chemically active amino acids and many conjugation methods involve amino acids such as lysine, cysteine, tyrosine, as well as N-terminal amino group and C-terminal carboxyl group.^32^ To generate site-specific positioning of nanoparticles placement of the amino acids accessible for labelling is critical. For the protein superhelix, we made site mutations to create multiple lysine groups (Figure 1D, lysine nitrogen atoms highlighted with red spheres. Amino acid sequence in Supplementary Information). The newly-introduced lysine groups on the structure have a well-defined spacing of 15.4Å from the model and are located at the exterior face where are expected to be more accessible for labelling guest molecules. To facilitate the protein purification, a poly-His tag was added at the N-terminus followed by a TEV cleavage site and a two-Gly residue spacer (Figure S1). A codon optimized gene corresponding to the sequence was synthezised and cloned into a pET-21 vector. Protein production was carried out in BL21 competent E. coli with standard protein expression methods (Experimental section). Protein purification was performed using a His-tag affinity column filled with Ni-NTA resin.

**Figure 2:**
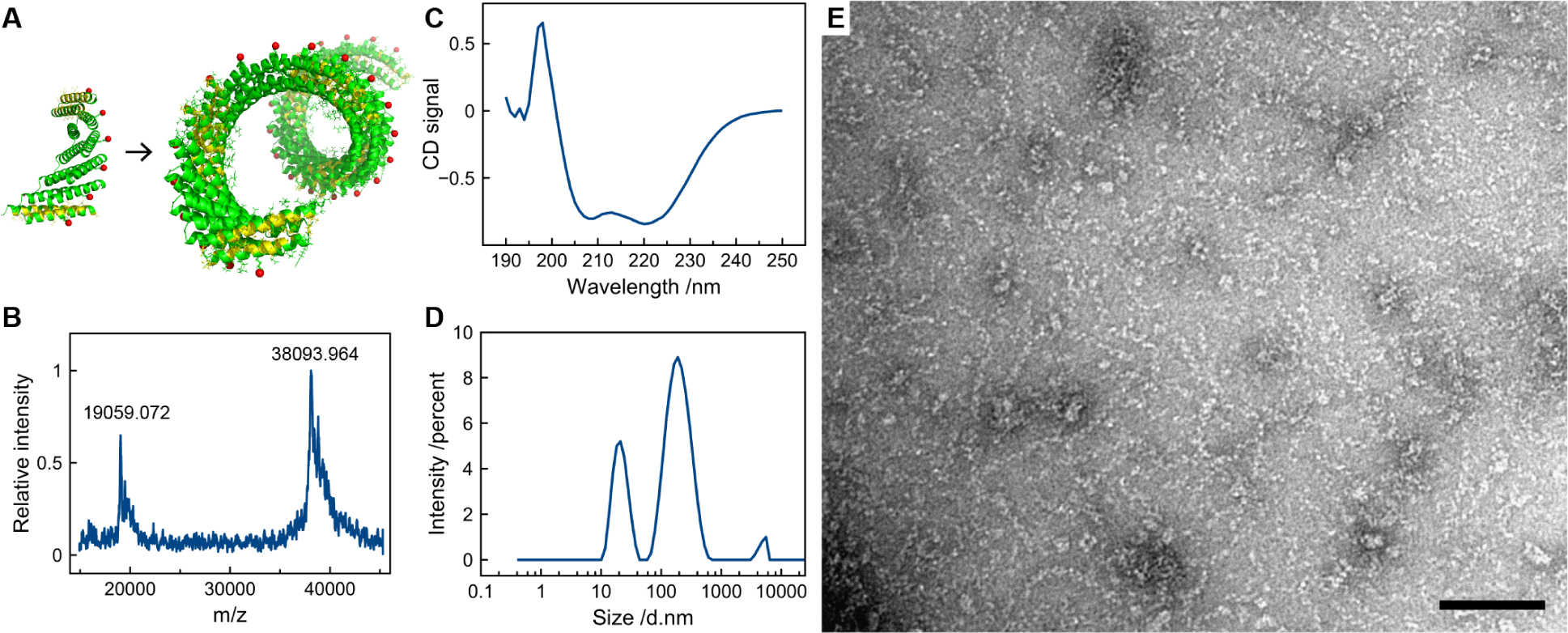
The characterization of protein superhelix using different techniques. A, the scheme of the superhelix formation from a chiral monomer. Red spheres highlight the lysine nitrogen atoms. B, MALDI-TOF mass spectrum shows the protein monomer peak at 38K and the double icons peak at 19 K. C, Circular dichroism spectra confirmed the successful folding of alpha helix in monomer. D, Dynamic light scattering measurement showed a concentration dependent fibre sizes and scattering peaks at 20nm, 200 nm, and 600nm indicating large structures formed. E, Negatively stained transmission electron microscope images showed the helical protein fibre. Scale bar: 100 nm.

We first characterized the protein with MALDI-TOF mass spectroscopy. As shown in Figure 2B, mass spectrometry confirmed that monomers have the expected mass of 38237g/mol. Circular dichroism spectroscopy (CD) was characteristic of alpha-helical protein with two peaks at 208 and 222 nm indicating the successful folding of the monomer. Dynamic light scattering (DLS) measurements showed scattering peaks in the range of large size distributions, spanning from a few tens of nanometers to micrometers, as well as concentration dependence over sequential dilutions (Figure S2). Negatively stained transmission electron microscopy (TEM) showed the formation of one dimensional helical fibres (100 µM monomer concentration). Further TEM images analysis showed that the fibre has a diameter of about 6.5 nm and a pitch around 8.9 nm, which agrees well with the expectation from the structural model (5.1 nm diameter with 13.1 nm helical pitch). We also observed the fibres have different lengths from several 10 up to 200 nm. Size exclusive chromatography (SEC) showed multiple peaks (Figure S3) which is consistent with protein fibres of various lengths as observed from TEM. However, non-specific interactions from the tremendous amount of hydrophobic residues (highlighted in yellow, Figure 1C, Figure 2A) at the ends of protein helical fibres could potentially prolong the retention time during SEC by adsorbing to stationary phase and complicate the analysis of the chromatography results.

We selected gold nanoparticles to demonstrate the templated assembly due to their commercial availability and well-studied surface functionalization. To introduce DNA, the dual linker GMBS (N-*γ*-maleimidobutyryl-oxysuccinimide ester) was conjugated to protein amino groups in 100 mM, pH 7.2 HEPES buffer (Figure 3A). Excessive amount of GMBS was filtered out using 100 kDa MWCO Amicon filter. DTT (Dithiothreitol) treated thiolated DNA (thiol-19A) was then added to attach to the maleimide group of GMBS on protein. After purification by filtration with three times washing, DNA-protein conjugates were obtained. Absorption measurement shows that after DNA conjugation, a new peak appears at 260 nm, a characteristic absorption of DNA (Figure 3B). The thiol-polyT (19T) capped 10 nm gold nanoparticles were mixed with polyA-protein conjugates in a molar ratio of 2:1, using non-DNA modified protein as control. Without further purification, as shown in TEM images (Figure 3C and 3D), the gold nanoparticles with non-DNA modified protein showed no assembly behaviour. In contrast, for the gold nanoparticles mixed with DNA-protein conjugates, gold nanoparticles were organized into short chains. The chain lengths are in a good agreement with the length of protein superhelix fibres. Due to the small diameter and short pitch of protein helix, however, 10 nm gold nanoparticles assemblies were not able to reflect the superhelical feature.

**Figure 3.**
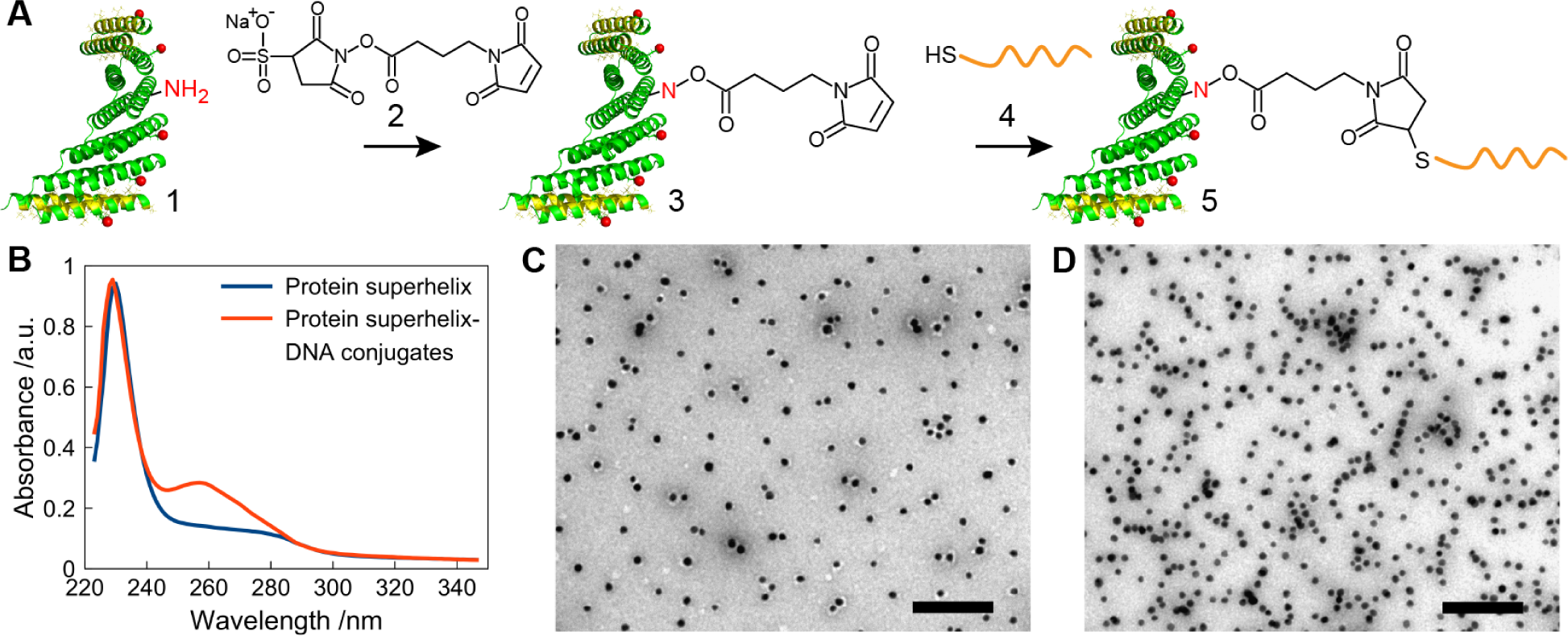
DNA-mediated gold nanoparticles assembly on protein superhelical structures. A, Stepwise reaction to conjugate thiolated DNA to protein primary amine groups. 1) protein superhelix, 2) hetero-dual linker GMBS (N-γ-maleimidobutyryl-oxysuccinimide ester), 3) protein superhelix conjugated with GMBS, 4) thiolated DNA oligonucleotide with sequence 19A, 5) protein-DNA conjugate. B, UV absorption of protein superhelix (blue curve) and protein-DNA conjugate (red curve). C&D, transmission electron microscopy image of gold nanoparticles functionalized with thiolated 19T mixed with protein superhelix without (C) and with (D) DNA modification. Scale bars: 100 nm.

To conclude, we presented DNA-mediated gold nanoparticles self-assembly on a protein superhelix formed from *de novo* designed alphahelical building blocks. The presented methodology has several benefits. The orthogonal DNA sequences design permits addressable assembly even for different types of nanocomponents simultaneously. At the same time, nanoparticles aggregations will be minimized as a result of surface functionalization with DNA. The rapid development of methods in protein design have enabled engineering of complex protein templates with well-defined placement of functional residues. In particular, simple repeated protein assemblies^18^ from individual monomers have demonstrated molecular-scale control combined with long-range assembly capabilities. Sophisticated assemblies can be generated through self-assembly of homomeric building blocks with precisely-defined interactions, topologies, and directionality. The integration of complex protein templates design and DNA-mediated conjugation will enable the organization of various nanocomponents on protein structures and pave the way for the systematic exploration of the physical coupling events in self-assembled nanomaterials.

## Experimental

### Protein expression

The monomer (6 pairs of repeated alpha-helix) was cloned into the pET-21a(+) vector (Genscript) containing an N-terminal 6*His-tag. BL21 E. coli bacterial strains were grown on LB agar plates casted with ampicillin and chloramphenicol in LB broth (1% Tryptone, 1% NaCl and 0.5% Yeast extract). Large scale protein expression was induced by 1 mM IPTG in LB broth, with antibiotics ampicillin and chloramphenicol, from a single colony. The incubation condition in the protocol is 37 °C with shaking speed 200 rpm unless otherwise stated.

In detail, 100 ng of DNA was added to 20 *µ*l BL21 E. coli cells and the mix was incubated on ice for 60 mins, followed by heat shock at 42 °C for 40 seconds resulting in the transformation of plasmid DNA into E. coli. The cell solution was then added to 1 ml SOC (2% Tryptone, 0.5% Yeast extract, 10 mM NaCl, 2.5 mM KCl, 10 mM MgCl_2_, 20 mM glucose) medium, followed by an incubation for 30 - 60 mins. 30 *µ*l cell solution were plated on LB-agar plate and incubated overnight to select the transformants. Bacterial colony plates were stored at 4 °C for future use.

Overnight starter cultures consisted of a stab from an isolated colony into 5 ml of LB medium containing 100 *µ*g/ml ampicillin and 34 *µ*g/ml chloramphenicol for antibiotic selection. This starter culture was used to inoculate a 400 ml culture at an initial OD of 0.1 using the same concentration of antibiotics. Induction of protein expression was performed by the addition of 1mM IPTG when the cells reached an OD of 0.5 while maintaining growth at 37°C. Following a 4 hr induction, cells were harvested *via* centrifugation at 6000 rpm for 12 min. Approximately 7.4 g of cell paste was recovered and resuspended in 20 ml of lysis buffer (50 mM Tris, pH 7.5, 150 mM NaCl, and 2 Roche complete protease inhibitor cocktail tablets). Cells were lysed on ice *via* sonication with 8 total cycles. Each cycle consisted of 30 seconds of sonication, alternating 1 sec on and 1 sec off, followed by a 30 sec interval for cooling. Benzonase nuclease (250 U) was added to the lysate, followed by 15 mins incubation at RT. Cellular debris was pelleted by centrifugation at 27000 x g for 30 min. SDS PAGE was used to assess the protein product and purity.

### Purification using his-tag affinity column

His-tag affinity column was made by adding 2 ml Ni-NTA resin (ThermoFisher) to a 20 ml column. Before loading the protein, the column was washed with 2 ml elution buffer (50 mM Tris, 150 mM NaCl, pH 7.5, 500 mM Imidazole), followed by a rinsing using 20 ml washing buffer (50 mM Tris, 150 mM NaCl, pH 7.5). Protein solution after centrifugation was then loaded to the column with a gravity flow speed of about 1 ml/min, followed by a washing using 20 ml washing buffer (50 mM Tris, 150 mM NaCl, pH 7.5, 20 mM imidazole). Target protein were washed out from his-tag beads using about 6 ml elution buffer (50 mM Tris, 150 mM NaCl, pH 7.5, 500 mM Imidazole) and collected in 0.5 ml aliquots.

### Instrumental information

Mass spectrum was recorded using MALDI-TOF MS, Applied Biosystems in the linear configuration. CD spectra were recorded in a quartz cuvette with 1 mm path length on Jasco spectropolarimeter with operation temperature at 20°C. Dynamic light scattering measurement was carried out on Malvern Zetasizer using disposable plastic micro cuvettes. For transmission electron microscopy, protein sample was deposited on carbon-coated copper grids and negatively stained using 1% uranyl acetate for 10 sec for inspection on a FEI Tecnai Spirit BioTWIN transmission electron microscope. The dimensions of the fiber and the helical geometry was measured with the software package ImageJ (http://imagej.nih.gov/ij/). 59 diameters and 21 helical pitches at selected domains were manfully marked with ImageJ and averaged as shown in the main text.

## Acknowledgement

The authors thank Lina Gefors for the assistance on TEM imaging and Katja Bernfur for the help of using Mass spectroscopy. The authors thank Dr. Ryan Oliver for helpful discussions and proofreading. Tao Zhang was supported by Carl Tryggers fellowship.

## Graphical TOC Entry

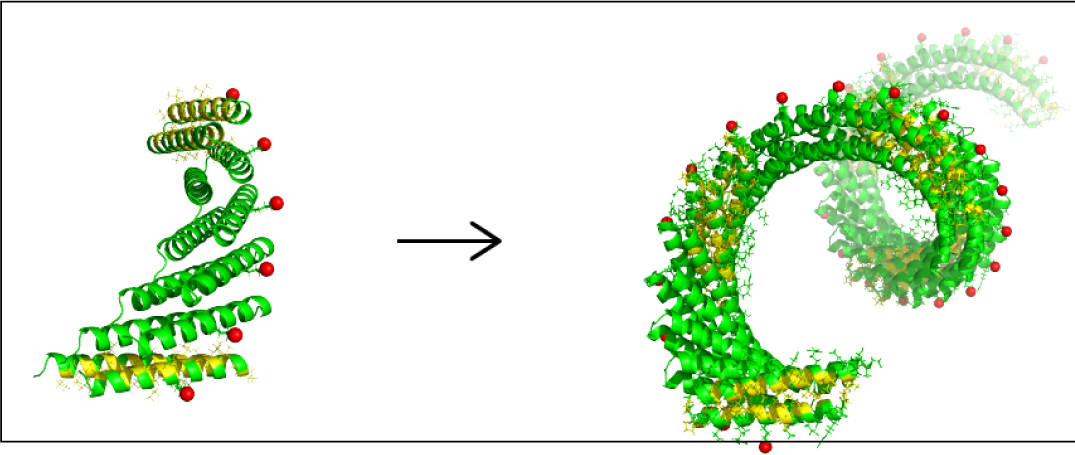

